# Effect of air pockets in drug delivery in jet injections

**DOI:** 10.1101/2021.02.02.429451

**Authors:** Pankaj Rohilla, Emil Khusnatdinov, Jeremy Marston

## Abstract

Needle-free jet injections are actuated by a pressure impulse that can be delivered by different mechanisms, and the resultant jets are 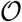(10^2^) m/s. Here, we report on the effect of entrapped air bubbles since filling procedures for pre-filled ampoules can induce bubbles, especially for viscous fluids. We use spring-piston devices as the principal actuation mechanism and vary both the location and size of the initial bubble. We find that the bubble location does have a statistically significant (*p* < 0.05) effect on the jet exit speed, based upon the volumetric flow rate. However, we reveal subtle features such as intermittent atomization when the gas pockets pass through the orifice and de-pressurize, which leads to spray formation and a temporary increase in jet dispersion, both of which can lead to product loss during an injection. These results have implications for the development of prefilled ampoules for jet injection applications.

## INTRODUCTION

Conventional oral administration of drugs is limited by low bioavailibility caused by the acidic environment in the stomach and restrictive intestinal epithelium [1]. The most common alternative to oral drug delivery are hypodermic needle syringes, which have numerous caveats including needle-stick injuries, cross-contamination, and needle-phobia [2, 3, 4] and potential limitation on injectability of viscous drugs. Thus, evidently there is a need for an effective needle-free technique for drug administration. Needle-free jet injections are one of the alternative techniques where a high-speed narrow jet (diameter, *d_j_* ~100-200 μm) punctures the top layer of the skin *(i.e. stratum corneum)* and deposits the drug into the intra-dermal/subcutaneous/intramuscular tissue [5].

Hypodermic syringes and needle-free jet injectors are medical devices which need an ampoule or barrel to store medication or biological drugs before injection. These ampoules are either supplied prefilled or can be loaded manually. One of the advantages of prefilled ampoules is the low variability in the amount of drug without any air pocket [6]. However, air pockets can form within the liquid inside the ampoules during loading or transportation, which can further be amplified by the viscosity effects of the contained fluid. Such air pockets can affect the volume and other variables related to drug delivery. The effect of air pockets on the efficacy of drug delivery is still largely unknown. This demands a systematic parametric study to further the current understanding.

In general, the presence of air pockets within a drug is either undesirable or could be harnessed in several medical applications[7, 8, 9, 10, 11]. Ultrasound imaging [12, 13], lithotripsy [14], controlled cavitation in gene delivery [13] and laser-induced jet injection [15, 16, 17] are some of the applications where energy generated due to the formation and collapse of air pockets can be utilized. On the other hand, the phenomenon of collapsing air pocket also generates microjets and shock waves, which can rupture red blood cells in artificial heart valves and can be detrimental to soft tissue [18, 19, 20].

Introducing air bubbles *(0.1-0.2 ml)* before drawing medication into a syringe was commonplace practice among clinicians and nurses in the United States before the widespread use of disposable syringes in the 1960s [6, 21]. These air bubbles were used to correct the medication dosage to account for the dead volume in the needle hub. Moreover, air pockets when injected with the medication were thought to prevent the seepage or backflow of the medication into the subcutaneous layer through the needle tracks [22]. Multiple studies showed contradictory results to this hypothesis [23, 24]. Moreover, air bubbles occupy the available volume for the medication which cause dosage inaccuracy [25]. Thus, it is recommended to avoid air bubbles in syringe injections [6, 21, 25].

In jet injection, air bubbles can affect the coherency and continuity of the jet stream in addition to dosage errors. This phenomenon intensifies with an increase in plunger speed. Thus, in the context of jet injectors where the jet speeds can be 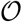(10^2^) m/s, it is important to study the effect of air pockets in the nozzle on drug delivery via jet injection. Although researchers have studied the effect of cavitation in the storage and injection of therapeutics in the past [7, 8, 26, 27, 28, 29, 30, 31, 32], there is a lack of detailed study to understand the effect of air pockets in intradermal injections via jet injectors.

Here, we study the effect of air pockets present at different locations within the nozzle cartridge on the hydrodynamics and efficacy of drug delivery via jet injection. The physical properties of the liquid and nozzle dimensions were kept constant. We studied the atomization of the bubbles within the liquid and the dispersion of the jet when bubbles exit through the orifice. In addition, ex vivo studies were conducted on porcine skin to understand the effect of air pockets on the delivery efficiency.

## MATERIAL AND METHODS

A spring-based jet injector *(Bioject ID pen)* was used in the experiments, which is described in detail in other works [33, 34, 35], but comprises a stiff spring piston with a cartridge with an orifice of diameter, *d_0_~157μm*. DI water was filled in a transparent nozzle cartridge with a volume capacity of 0.11 ml. Air pockets were introduced at different locations inside the nozzle cartridge *(as shown in figure 1(a))* using 1 ml luer-lock syringe with 24-gauge needle. Furthermore, the plunger was carefully replaced to avoid the escape of air pockets via the orifice exit. We have used five different locations within the nozzle cartridge to understand their effect on the bubble dynamics with time. These locations were selected on the basis of their probability of occurrence. For location *L_1_*, an air pocket was introduced at or in close proximity of the plunger tip. It should also be noted that the bubble locations, *L_2_* and *L_3_* are isotropic radially.

**Figure 1:**
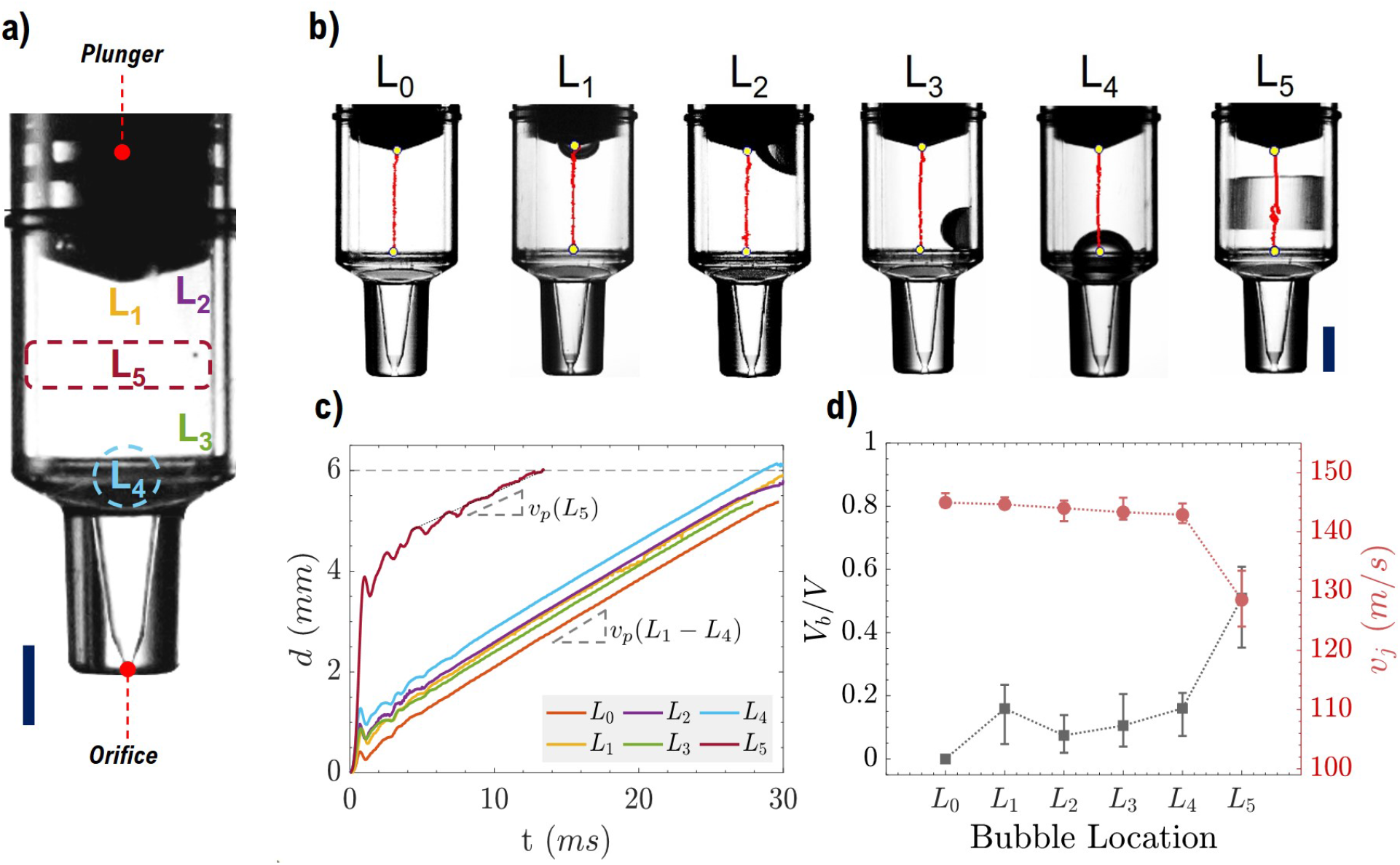
Air pockets in a nozzle cartridge. **(a)** Five different locations of air pockets inside a nozzle cartridge, **(b)** trajectory (in *red*) of plunger motion inside the nozzle for different locations of air pockets, **(c)** plunger-displacement with time for a jet injection with and without air pockets, and **(d)** volume fraction *(V_b_/V)* and jet speed *(υ_j_)* for different locations of air pockets. *Scale bar represents 3 mm (a,b)*.

To capture the bubble and jet dynamics, a high-speed video camera *(Phantom V711, Vision Research Ltd.)* was used at frame rates in a range of 10,000-30,000 frames per second. The tip of the plunger was tracked to get the plunger displacement *(figure 1(b))* with time using Photron FASTCAM Analysis *(PFA ver. 1.4.3.0)* software. Plunger speed was then estimated from the slope of the displacement-time plot after the initial ringing phase, e.g. *t >* 5 ms *(figure 1(c))*. Furthermore, we estimated the jet speed at the orifice exit, *υ_j_ (figure 1(d))* from plunger speed using mass conservation, assuming the liquid to be incompressible at the experimental conditions.

A single side-view perspective was sufficient to track the plunger tip. However, to understand the bubble dynamics and for qualitative analysis of bubbles within the nozzle during a jet injection, we employed two cameras *(second camera: Phantom Miro 311, Vision Research Ltd.)* with orthogonal views. In addition, using extension tubes with Nikon micro-nikkor 60 mm lenses, we achieved typical effective pixel sizes in a range of 10-30 *μm/px*.

Volume fraction was used to quantify the size of the air pockets, which can be defined as a ratio of the volume occupied by an air pocket (V_b_) and the total liquid volume inside a nozzle cartridge without any air pocket *(V)*. We measured the volume fraction occupied by air pockets at different locations by weighing the liquid-filled nozzle with and without air pockets. Figure 1(d) shows the volume fraction of air pockets introduced at different locations inside the nozzle.

To measure the force profile during the jet injection, a miniature load button cell *(Futek-LLB 130, 50 lb, FSH03380)* was placed at a distance of 2 mm away from the orifice exit of the nozzle to avoid contact with the nozzle during jet injection. Force profiles of jet injection were recorded at a sample rate of 4,800 Hz.

To conduct ex vivo studies, porcine skin was used as a skin model for human skin. Porcine skin patches *(thickness ~3-5 mm)* were harvested from the side regions of Yorkshire-Cross pigs *(age: 13 weeks)*, euthanized in the department of Animal Sciences *(Texas Tech University*). These skin patches were kept in a −80°C freezer, but thawed to room temperature before jet injection. We cut skin across the center of the injection site to visualize the dispersion of the liquid after jet injection. Trypan Blue *(Sigma Aldrich)* was used as a dye (*1 mg/ml*) in DI water to aid in the visualization of skin blebs, and a custom Matlab script was used to estimate the dimensions of skin blebs. One must note that although fresh porcine skin is a close model to human skin at high humidity conditions [36], we used skin after a single freeze-thaw cycle.

We performed one-way ANOVA tests to check the statistical significance of various parameters used in the study with a significance level of α = 0.05.

## RESULTS AND DISCUSSIONS

### Bubble Dynamics

Air pocket collapses with the inertial impact of the plunger, atomizing into multiple microbubbles. We captured the qualitative behavior of these micro bubbles with time as the plunger pushes the liquid out via a narrow orifice. As shown in figure 1(c), plunger motion exhibits a ringing phase caused by the sudden impact of the spring piston on the plunger, followed by a nearly linear and stable motion. Average jet speeds were calculated from the slope of this linear region of the plunger displacement-time plot. Introduction of air pockets did not alter the jet speed *(υ_j_)* significantly which lied in a range of ~142-146 m/s except for the case of air pocket introduced at the center of the nozzle cartridge (*L_5_*).

Among the different locations of air pockets, *L_5_* showed a distinctive behavior. The plunger moved ~3.7 mm within a short period of 1.5 ms, thus resulting in a rapid collapse of the air pocket. The collapsing air pocket generates a pressure opposing the plunger motion, and the interplay of forces during this period is manifested by the recoiling trajectory of the plunger, as shown in the last frame of figure 1(b). The impulse is much stronger, and resistance in the early stages is much weaker, leading to significant over-pressure (noted by a force increase of ~33%). This resulted in a prolonged recoil/ringing phase. Due to this extended ringing phase, we calculated the average jet speed by fitting a line after this phase to the plunger displacement-time plot. For *L_5_*, average jet speed was lower (128.5±3.8 m/s). Another important characteristic of air pockets introduced at different locations inside the nozzle was the volume fraction of the air pocket, which was nearly half at location *L_5_*. In the case of other locations, the air pockets occupied ~ 20% of the total available volume.

Figure 2 shows a montage of frames summarizing the bubble dynamics for air pockets placed at different locations. When the plunger is triggered, the fluid pressure in the cartridge increases rapidly up to 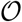(10 MPa), causing the air pocket to collapse and disintegrate into microbubbles. The trajectory of such microbubbles largely depends on the pressure gradient, which further depends on the flow disturbance caused by the bubbles collapsing under high pressure. Although numerous studies have been conducted to understand the behavior of bubbles with inertial impact [29, 30, 37], the underlying physics behind the bubble dynamics may vary on the basis of applications.

**Figure 2:**
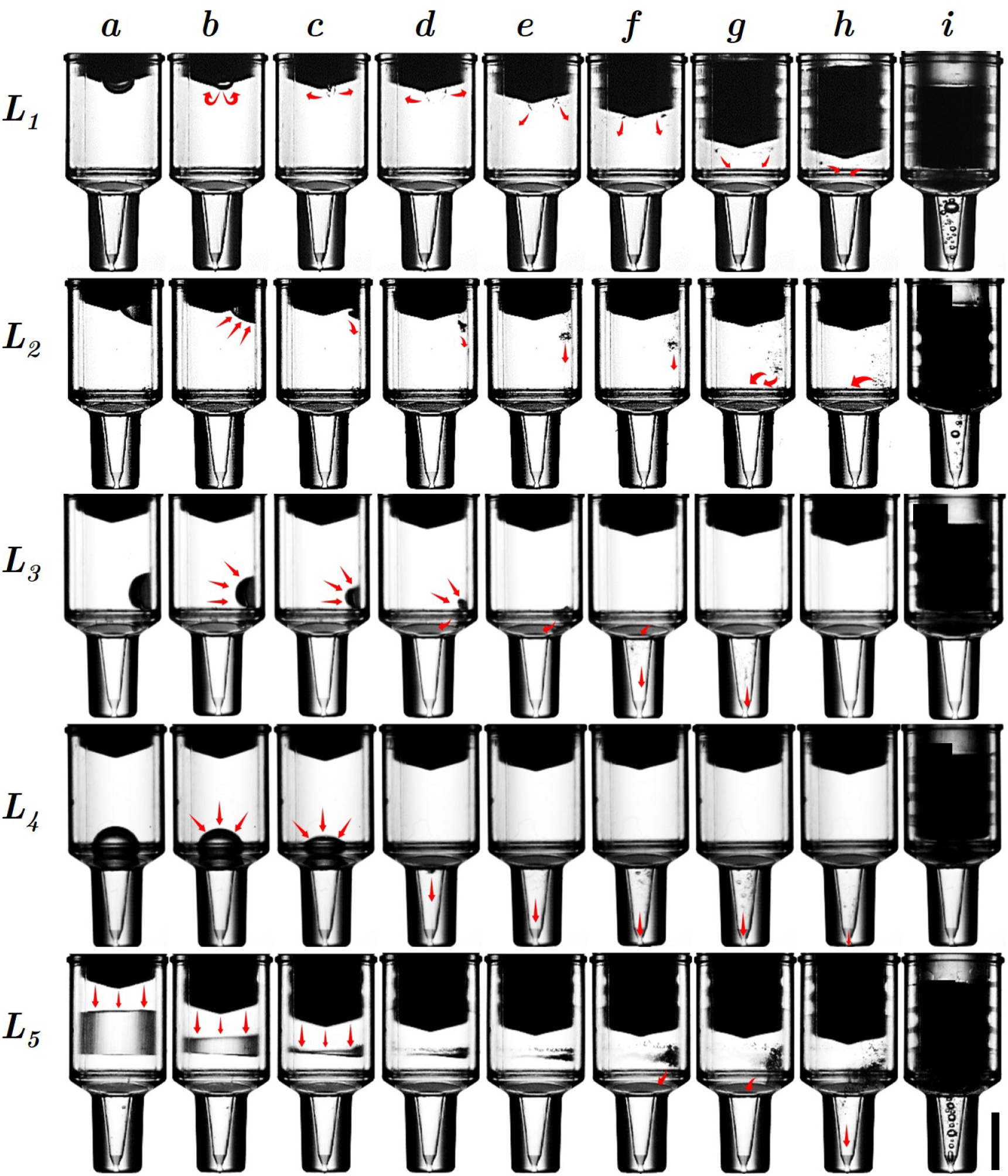
Snapshots of bubble dynamics for different locations of air pockets inside the nozzle cartridge. Snapshots of bubble dynamics corresponding to different time frames for location *L_1_ ≡(a – i)* ≡ [0, 0.1, 0.2, 0.27, 0.37, 1.5, 2.27, 5.73, 61.07] ms; **for location** *L_2_(a – i) ≡* [0, 0.2, 0.3, 0.5, 1.0, 2.9, 4.57, 6.23, 61.77 ms; **for location** *L_3_(a – i)v ≡* [0, 0.1, 0.2, 0.27, 0.37, 1.5, 2.27, 5.73, 61.07] ms; **for location** L4(α – i) ≡ [0, 0.67, 0.167, 0.43, 0.67, 1.47, 2.23, 3.5, 64.67] ms; **for location** *L_5_(a – i) ≡* [0, 0.5, 0.6, 0.67, 0.73, 1.1, 1.83, 5.3, 30.33] ms.

The life cycle of microbubbles depends on the location where the air pocket was introduced initially. For air pockets present near the outlet (*L_3_* and *L_4_*), microbubbles exit the nozzle at an early stage whereas for air pockets introduced farther upstream, not all of these bubbles exit; it can be noticed from *i^th^* frames in figure 2 that microbubbles were present in the tapered section of the nozzle at the end of injection for air pocket introduced at locations *L_1_, L_2_* and *L_5_*.

Figure 3 shows orthogonal views of microbubbles inside the nozzle at same instant (t = 5.5 ms). Different views shown in figure 3 yield different bubble size distributions at the same instant of time due to the lensing effects of the nozzle geometry. Thus, to avoid the erroneous measurement, we did not quantify the bubble size distribution with time.

**Figure 3:**
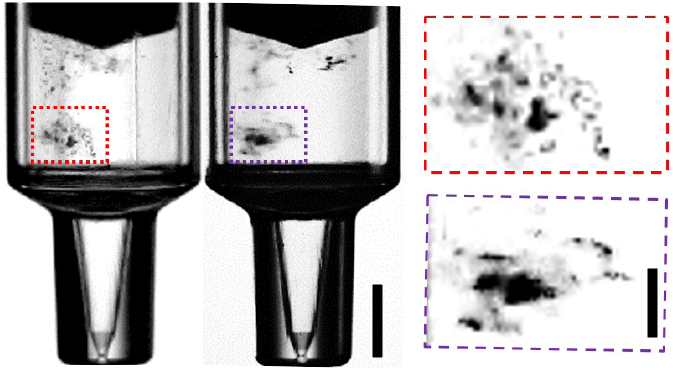
Orthogonal views of nozzle cartridge with microbubbles inside the liquid corresponding to time, *t* = 5.5 ms for air pocket introduced at location L_2_. Scale bar represents 3 mm. **Note:** A vertical line on the left frame is an artifact of the nozzle and is not related to the contained fluid or air pockets.

Assuming the air bubble to be an ideal gas, we can estimate the volume of the wedge-shaped bubble cloud formed after the collapse of an air pocket at location L_5_, using ideal gas law (*PV* = *nRT* = *const.):*

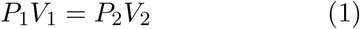

Where *P_1_* and *P_2_* are atmospheric pressure (~10^5^ P_a_) and pressure inside the bubble (~5×10^7^ P_a_, using peak force of *F*\* ~1 N), respectively. *V_1_* and *V_2_* are initial volume of air pocket (~55 μL) and the volume of the wedge-shape bubble cloud after the collapse, respectively. Minimum height of the bubble cloud *(h_min_)* was estimated using:

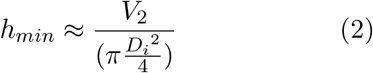

Where *D_i_* is the inner diameter of the nozzle barrel.

A slight asymmetry in the plunger causes a wedge-shaped air volume that disintegrates from left to right in the image sequence *L_5_(a-i)*, therefore, we can expect a size distribution with *r_min_* « 10 μm and *r_max_* » 10 μm, evidenced by zoomed images where bubbles with size of 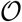(100 μm) can be easily seen. Moreover, many bubbles with a size of the order of 1 pixel or less were also observed.

### Force measurements

To measure the impact force, liquid jets were impinged on a load cell placed at a distance of 2 mm away from the orifice exit. Figure 4 shows force profiles of jets with air pockets present at different locations. The time duration of the jets for air pocket locations *L_1_, L_2_, L_3_, L_4_ and L_5_* were ~37.5 ms, ~34.8 ms, ~35.8 ms, ~33.5 ms, and ~16.9 ms, respectively. In the absence of an air pocket, the time duration of a jet was ~38 ms. It is noteworthy that the time duration of injection was lower in displacement-time plot *(figure 1(c))* due to the difficulties associated with tracking the plunger tip towards the end of injection.

**Figure 4:**
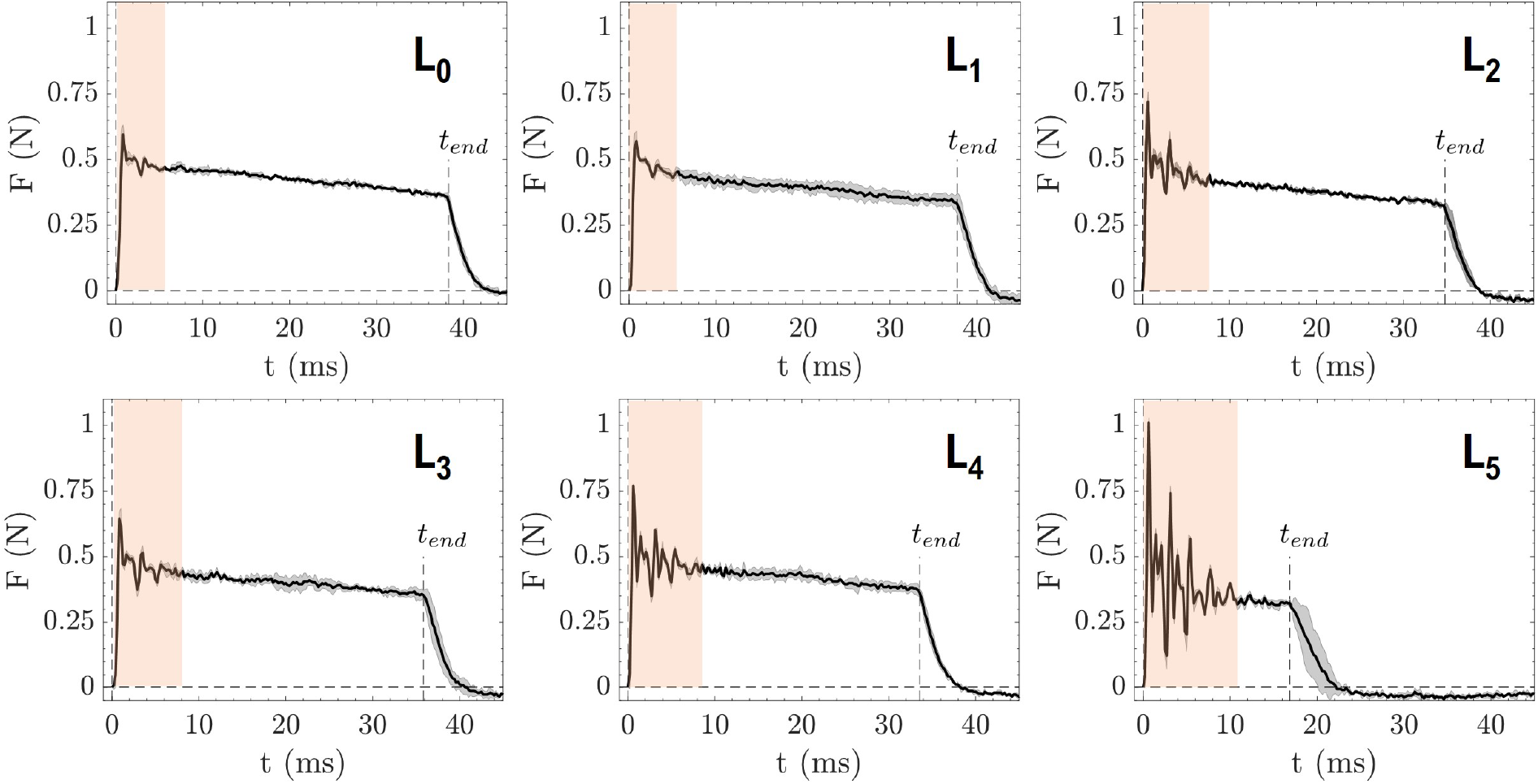
Impact force profiles for jet injections for different locations of air pockets. *L_0_* represents the jet injection without any air pocket inside the nozzle. Shaded region represents ringing phase during jet injection. *n=3*.

As the spring piston strikes the plunger, a ringing phase can be observed in force profiles in the initial stage (~10 ms) of liquid injection. In the absence of any air pockets, the peak force was ~0.63 N and a similar peak force was observed for air pockets present at locations *L_1_, L_2_, L_3_*, and *L_4_* with values of ~0.62 N, ~0.76 N, ~0.67 N, and ~0.78 N, respectively. However, *L_5_* exhibited a higher force of ~1.03 N, with an increase of 63%. After the peak force, the magnitude of the force profile was nearly the same for all cases with 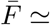 0.4 N (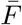 *= ρυ^2^ A_0_, where A_o_ is the cross-sectional area of orifice exit)* except the case of *L_5_* with lower magnitude of the measured force. Further implications of force profile and injection duration will be discussed in ex vivo studies.

### Effect of microbubbles on jet

As pressurized microbubbles exit the orifice, they undergo a rapid depressurization and expand, resulting in a spray-like jet (figure 5). The magnitude of the effect of microbubbles on the jet depends on the size and the number of bubbles exiting the nozzle orifice along the liquid jet.

**Figure 5:**
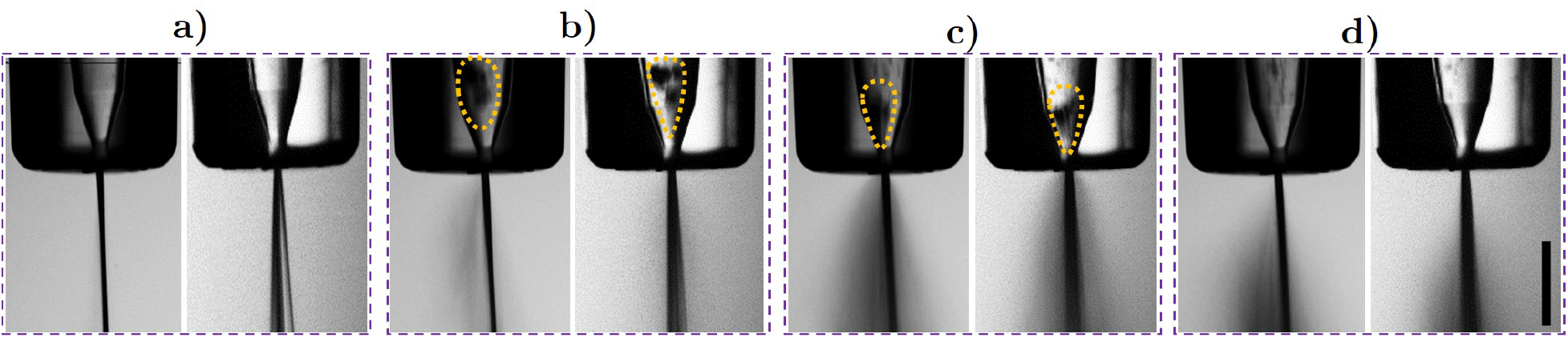
Effect of bubbles on the jet coherency with orthogonal views. (a) liquid jet without any bubbles (*t* = *1.4 ms*), (b) bubble cloud in tapered section before exit (*t* = *9.9 ms*), (c) spray-like jet formation as the bubbles exit with the jet (*t* = *10 ms*), and (d) diminishing atomization of jet as bubble clouds exits the tapered section (*t* = *10.1 ms)*. *Scale bar: 2 mm*

Figure 5 shows the effect of a bubble cloud exiting the nozzle on the liquid jet with orthogonal views captured using two high-speed cameras at a frame rate of 10,000 fps. The liquid jet free of any effect of bubbles is shown in figure 5(a), where orthogonal views show that the jet looks coherent from one side and slightly dispersed from the other side. In the next frame (figure 5(b)), a cloud of air pockets *(highlighted in dashed yellow outlines)* appeared in the tapered section of the nozzle with spray-like jets as the bubbles exit with the jet. As more bubble cloud exits the nozzle, the jet dispersion grows as shown in figure 5(c). With the majority of air pockets exiting the orifice exit, the remnant effects of the bubbles on the liquid jet can be seen in figure 5(d), but the jet will resume the steady stream upon clearance of air bubbles.

### Ex vivo studies

Dyed water was injected into the porcine skin to understand the effect of air pockets on the percentage delivery. Porcine skins were thawed to room temperature before administering injections. The thickness of the dermis layer of porcine skin harvested from different pigs (hence, different colors in figure 6) was within 3 mm. The effect of force exerted by the jet injector *(loading)* in the normal and axial direction during injection has been reported in an earlier study [35], and therefore we used a recommended normal load of 1 kg for the maximum delivery efficiency at which the injection was actuated.

**Figure 6:**
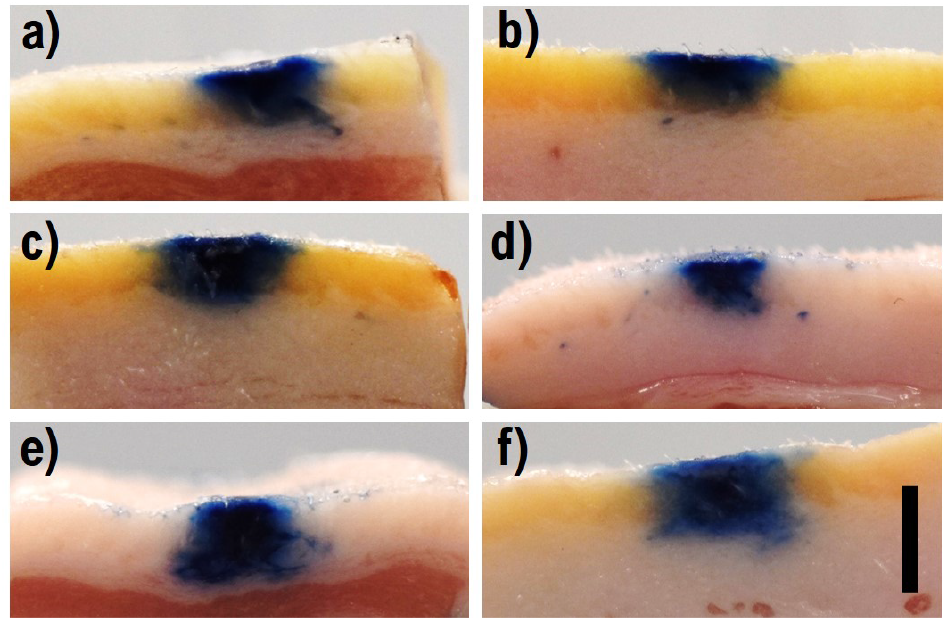
Cross section view of skin blebs for: **(a)** no air pockets, **(b)** air pocket at location *L_1_*, **(c)** air pocket at location *L_2_*, **(d)** air pocket at location *L_3_*, **(e)** air pocket at location *L_4_* and **(f)** air pocket at location *L_5_*. Scale bar represents 5 mm.

After impingement, the liquid jet creates a hole in the skin through which liquid propagates inside the skin. The stiff poro-elastic structure of the dermal tissue resists the liquid inflow inside the skin, thus, limits the amount of liquid that can be delivered with a single injection. To visualize the dispersion of liquid inside skin after jet injection, the skins were cut across the point of injection. Figure 6 shows cross-sections of skin blebs corresponding to the different location of air pockets.

Figure 7 shows the effect of the location of the air pocket on the delivery efficiency and the dimensions of the bleb formed inside the skin after jet injection. Delivery efficiency (η = (*m–m_r_*) × 100/m) was measured from the weight of liquid rejected at the top of the skin after jet injection *(m_r_)* and total available volume in nozzle cartridge *(m)*. Whatman filter papers were used to absorb the rejected liquid on the top of skin.

**Figure 7:**
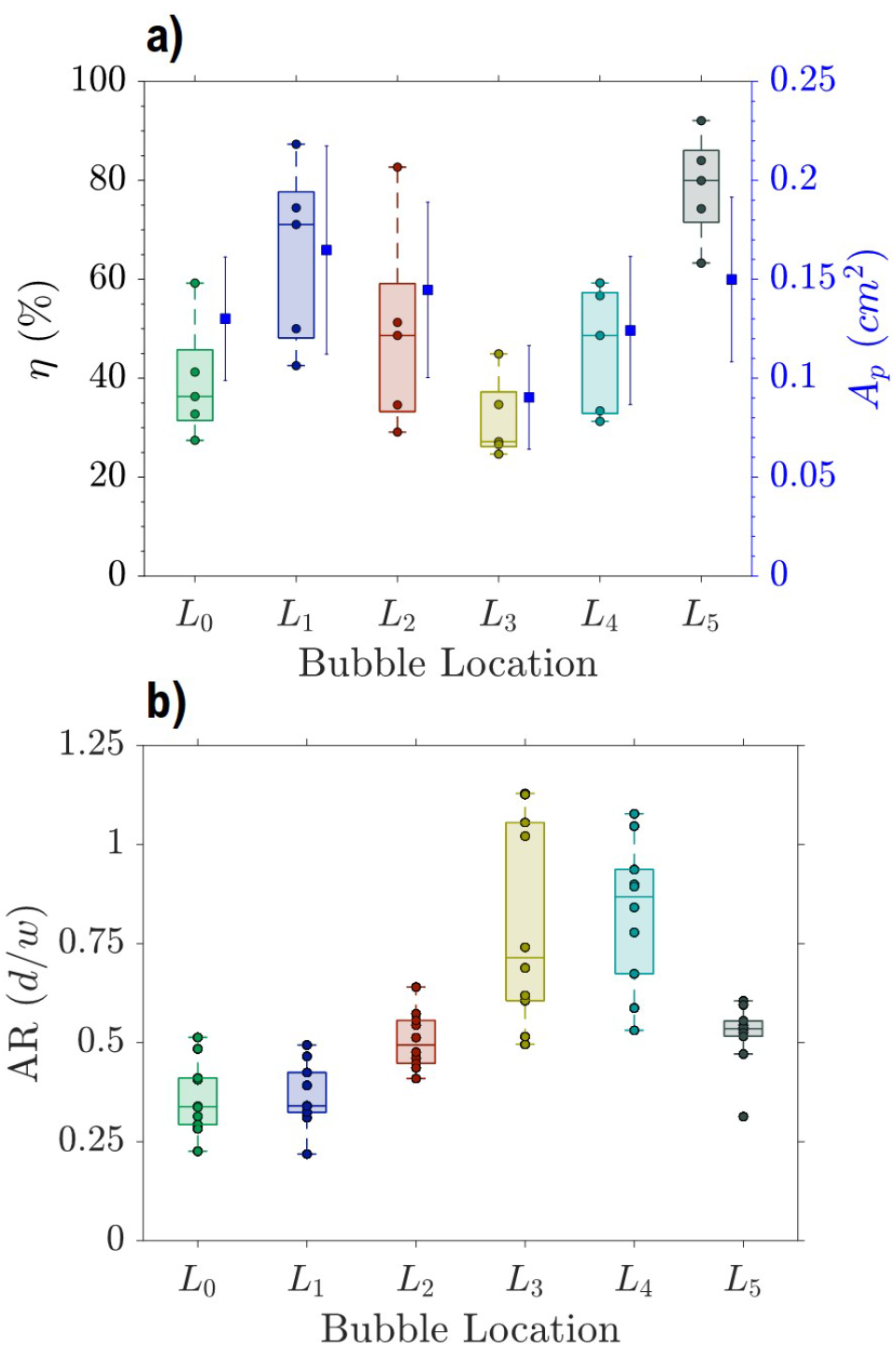
Effect of air pockets on effectiveness of drug delivery. **(a)** delivery efficiency and projected area of bleb cross-sections for jet injections with air pockets at different locations *(n=5)* and **(b)** aspect ratio of the blebs (*AR* = d/w), where *d* and *w* are depth and width of skin bleb respectively *(n=10*).

Presence of air pockets at locations *L_1_, L_2_*, and *L_5_* showed higher percentage delivery of liquid inside the skin. Effect of varying the location of air pockets on the delivery efficiency of dyed water was significant (p < 0.05). It is noteworthy that the delivery efficiency was affected by the confluence of changing location and available liquid volume inside the nozzle. As discussed earlier, the volume of liquid inside the nozzle changes with introduction of air pockets inside the nozzle. Introducing an air pocket at location *L_1_* showed higher percentage delivery than the case of no air pockets for nearly the same force profile. This increase in the percentage delivery could be done due to the reduced liquid volume to be injected inside the skin.

As shown in earlier studies, higher percentage delivery was obtained for jet injection of liquid in lower volumes [35]. It was hypothesized that there is a limit of liquid volume which can be injected into the skin without significant rejection. Here, with the introduction of air pockets, the available liquid volume to be injected was lower for locations *L_1_* and *L_5_*, resulting in higher efficiency.

In addition, the percentage delivery was nearly the same for location *L_3_* and *L_4_* as compared to control case of L_0_. Bubbles exit early with a jet for air pockets present at locations *L_3_* and *L_4_*. As the high-speed liquid jet penetrates skin, it creates a channel to facilitate the further delivery of incoming liquid. Any disturbance in the jet shape in the initial phase could have an adverse effect on the channel formation for liquid propagation, thus resulting in lower percentage delivery and narrow dispersion patterns, as observed for *L_3_* and *L_4_*. Higher peak force also helps in increasing percentage delivery. In the case of air pocket introduced at location *L_5_*, higher percentage delivery was observed; partly due to high impact force (~1.03 N) and lower available volume of liquid to be injected (~0.044 ml).

The effect of location of air pockets was significant on the projected area (*p <* 0.05) and the aspect ratio of skin blebs (*p <* 0.05). Projected area showed nearly the same trend as of delivery efficiency. In case of injection with an air pocket present near the exit (*L_3_* and *L_4_*), skin blebs showed large intrasample variation in their shape and their dimensions.

## CONCLUSIONS

We investigated the effect of introducing air pockets at various locations within the liquid contained in a nozzle cartridge for jet injectors. Air pocket collapses and forms microbubbles which affects the shape of the liquid jet. Air pockets at different locations affected the peak force during the initial ringing phase. Furthermore, ex vivo studies conducted on porcine skin showed nearly the same or higher percentage delivery of dyed water inside skin. We conclude that the high percentage delivery obtained for air pockets at different locations was due to lower available volume to be injected and higher peak force. Moreover, the projected area and aspect ratio showed a significant effect of various locations of air pockets in addition to a significant effect on percentage delivery. Although the introduction of air pockets helped in enhancing the delivery efficiency of the drug, air pockets should be avoided in the nozzle as it causes dosage inaccuracy which can alter the efficacy of the vaccine.

The measurement of the force profiles showed higher peak forces in the initial ringing phase due to hammer effect. Furthermore, ex vivo studies conducted on porcine skin showed similar or higher percentage delivery of dyed water inside the skin. We conclude that the high percentage delivery obtained for air pockets could be attributed to the lower available volume and higher peak force. In addition, the project area and aspect ratio showed a significant effect for various locations of air pockets in addition to a significant effect on percentage delivery. Although the introduction of air pockets helped in enhancing the delivery efficiency of the drug, air pockets should be avoided in the nozzle as it causes dosage inaccuracy which can alter the effect of the vaccine.

## ACKNOWLEDGMENTS

This work was financially supported by The National Science Foundation via award CBET-1749382.

## AUTHOR CONTRIBUTIONS

**P.R.** and **J.M.** designed the experiments. **E.K.** and **P.R.** conducted the experiments. **P. R.** analyzed the data and wrote the manuscript. **J.M.** reviewed and edited the manuscript.

