## Supplemental information document for "Effect of air pockets in drug delivery in jet injections"

### Supplementary Videos

All the supplementary videos were recorded at 10,000 frames per second and have play speed of 30 frames per second.
**Video 1:** Jet injection without any air pockets.
**Video 2:** Jet injection for an air pocket present at location $L_5$.
**Video 3:** Zoomed video showing the effect of micro-bubbles on the liquid jet for an air pocket at location $L_5$.
